# Gene-Gene interactions and pleiotropy in the brain nicotinic pathway associated with the heaviness and precocity of tobacco smoking among outpatients with multiple substance use disorders

**DOI:** 10.1101/782565

**Authors:** Romain Icick, Morgane Besson, El-Hadi Zerdazi, Nathalie Prince, Vanessa Bloch, Jean-Louis Laplanche, Philippe Faure, Frank Bellivier, Uwe Maskos, Florence Vorspan

**Author notes:** **Corresponding author:** Integrative Neurobiology of Cholinergic Systems, CNRS UMR 3571, Institut Pasteur, 25 rue du Dr Roux, Paris F-75015, France.

## Abstract

**Introduction:** Tobacco smoking is a major health burden worldwide, especially in populations suffering from other substance use disorders (SUDs). Several smoking phenotypes have been associated with single nucleotide polymorphisms (SNPs) of nicotinic acetylcholine receptors (nAChRs). Yet, little is known about the genetics of tobacco smoking in populations with other SUDs, particularly regarding gene-gene interactions and pleiotropy, which are likely involved in the polygenic architecture of SUDs. Thus, we undertook a candidate pathway association study of nAChR-related genes and smoking phenotypes in a sample of SUD patients.

**Methods:** 493 patients with genetically-verified Caucasian ancestry were characterized extensively regarding patterns of tobacco smoking, other SUDs, and 83 SNPs from the nicotinic pathway, encompassing all brain nAChR subunits and metabolic/chaperone/trafficking proteins. Single-SNP, gene-based and SNP × SNP interactions analyses were performed to investigate associations with relevant tobacco smoking phenotypes. This included Bayesian analyses to detect pleiotropy, and adjustment on clinical and sociodemographic confounders.

**Results:** After multiple adjustment, we found independent associations between *CHRNA3* rs8040868 and a higher number of cigarettes per day (CPD), and between *RIC3* rs11826236 and a lower age at smoking initiation. Two SNP × SNP interactions were associated with age at onset (AAO) of daily smoking. There was pleiotropy regarding three SNPs in *CHRNA3* (CPD, AAO daily smoking), *ACHE* (CPD, HSI) and *CHRNB4* (CPD, both AAOs).

**Discussion:** Despite limitations, the present study shows that the genetics of tobacco smoking in SUD patients are both distinct and partially shared across smoking phenotypes, and involve metabolic and chaperone effectors of the nicotinic pathway.

## Introduction

Tobacco smoking is responsible for six millions deaths per year worldwide, accounting for 12% of all deaths among adults aged ≥ 30 [1]. It is two to 13 times more prevalent in people with other substance use disorders (SUDs) than in control subjects [2], and SUD individuals present many risk factors for more severe patterns of tobacco smoking [3]. Consequently, tobacco-related diseases may account for 36-49% deaths of people who ever received inpatient treatment for other SUDs [4].

Tobacco smoking is a complex behaviour that encompasses several measurable phenotypes: smoking initiation and persistence, regular/dependent smoking, as well as cessation attempts, manifestation of withdrawal symptoms, treatment response and relapse. The main addictive component of tobacco, nicotine, binds nicotinic acetylcholine receptors (nAChRs), which are pentameric ligand-gated ion channels co-assembled from α and β subunits with both spatial and functional specificity in the brain [5]. Interestingly, smoking phenotypes have been repeatedly associated with frequent single nucleotide variants (called single nucleotide polymorphisms, SNPs) of nAChR subunit genes [6], mainly in the *CHRNA5-A3-B4* cluster on chromosome 15, which encodes the *α*5, *α*3 and β4 subunits. A non-synonymous *CHRNA5* rs16969968 – D398N SNP doubles the risk to develop nicotine dependence in homozygous carriers of the risk allele [6]. Other variants from the cluster, often in high linkage disequilibrium (LD) with each other and thus constituting several haplotypic blocks [6], modulate *CHRNA5* expression and the risk conferred by rs16969968 [7]. Most of the phenotypic components of tobacco smoking -especially age at onset (AAO) and the number of cigarettes smoked per day (CPD) – are strongly associated with the level of exposure to tobacco and, thus, to its morbidity. Yet, despite the very strong co-morbidity between tobacco smoking and other drug dependences, the influence of nAChR-related genetics of such tobacco smoking phenotypes in SUD patients has been insufficiently studied so far.

The biology of the nicotinic pathway supports the importance of studying gene-gene interaction to better consider its influence on complex traits such as tobacco smoking. Notably, essential to nAChR function are enzymes that metabolize acetylcholine (nAChR endogenous ligand) and trafficking/chaperone proteins [8–10] that have been overlooked to date in genetic studies focusing on nAChR genes. Interactions between *CHRNA4* and genes from the *Bdnf* pathway have been associated with the AAO [11] and the presence of nicotine dependence [12,13]. Finally, pleiotropy, *i.e.* simultaneous associations between a given gene or SNP with several smoking phenotypes is expected [14], as in most complex traits [15].

In this context, to fill the significant knowledge gap regarding the genetics of tobacco smoking in the context of other SUDs, we undertook a candidate pathway, within-case study to describe (i) the genetic architecture (broad nicotinic pathway) of relevant tobacco smoking phenotypes and (ii) gene-gene interactions further involved in these phenotypes, in a representative clinical sample of treatment-seeking outpatients with multiple SUD, who underwent extensive genotypic and phenotypic characterization.

## Methods

### Sample selection and clinical assessment

Patients > 18 years seeking treatment for any SUD other than nicotine as ascertained using DSM IV-TR criteria were consecutively recruited between April 2008 and July 2016 through two multicentric protocols. Both protocols and the present study were approved by the relevant Institutional Review Boards [CPP Ile-de-France IV and CEEI from the *Institut de la Santé et de la Recherche Médicale* (INSERM), IRB00003888 in July 2015, respectively]. All participants provided written informed consent for both the clinical and genetic assessments, and study records were continuously monitored by the local research administration (*Unité de Recherche Clinique*). The research was conducted in accordance with the Helsinki Declaration as revised in 1989. Eligible participants were excluded if they had severe cognitive impairment or insufficient mastery of the French language preventing misunderstanding of the study purposes and assessments, if they had no social insurance, and if they were under compulsory treatment. A unique standardized interview was conducted by trained investigators (Psychologists or M.Ds.). Smoking behaviour assessment included current smokers’ number of CPD and delay from wake-up to 1^st^ cigarette, which allowed for scoring *Heaviness of Smoking Index (HSI)* [a short and reliable tool for assessing nicotine dependence in current smokers (cut-off score ≥ 4) [16]] along with age at first cigarette use and age at daily cigarette smoking. The full study procedure is available as Supplementary Methods file 1.

### Biological sampling and genetic analyses

DNA was extracted from whole blood using a Maxwell 16 PROMEGA® extractor (Promega France, Charbonnières-les-Bains, France). Purity assessment followed the procedures described by the Centre National de Génomique, estimated on a NanoDrop® spectrophotometer by using Picogreen® assay, confirmed by Polymerase Chain Reaction before application on gel. Participants were genotyped using the Infinium PsychChip array (Illumina, San Diego, CA, USA) in two stages (2014 and 2017) by Integragen SA® (Evry, France) using the same pipeline. Genotype files were merged for the present study, keeping only bi-allelic variants common to both extractions.

### Genome-wide quality control and SNP selection

*PLINK* [17] was used for quality control (QC), based on a consensus procedure [18] (see flowchart, Figure 1), performed at the whole-sample (N=581) and genome-wide (566,932 SNPs) levels. Individuals of Caucasian ancestry and ≥2^nd^ degree relatives were identified by genotyping [identity-by-descent and principal component analysis by comparison with five 1000 genomes superpopulations (supplementary Figure 2]. QC and ancestry assessment left 493 individuals and 260,853 markers remaining. There were valid data regarding 433 (CPD), 370 (HSI), 470 (age at 1^st^ smoke) and 396 (AAO daily smoking) participants with a mean genotyping rate =99.831%. From the whole-genome DNA array, the study focused on 17 genes from the nicotinic pathway, encompassing those encoding human neuronal *α* and *β* nAChR subunits (*CHRNA2-7,9&10* and *CHRNB2-4*), cholinergic enzymes (acetylcholine esterase (*ACHE)* and choline acetyltransferase (*CHAT)*) and nAChR chaperone/trafficking proteins *LYNX1, LYNX2, RIC3, ZMYM6NB* (encoding *Nacho*), which provided 314 markers. HG19 was the reference human genome version. Eventually, 83 SNPs (minor allele frequency ≥5%) from 16 genes (no SNP from *ZMYM6NB*) of the nicotinic pathway remained after QC and marker selection. The full list of markers, their correspondence with Illumina® names and positions, gene length and coverage by the DNA array are listed in Supplementary Table 1 (final study markers are indicated with *).

**Figure 1:**
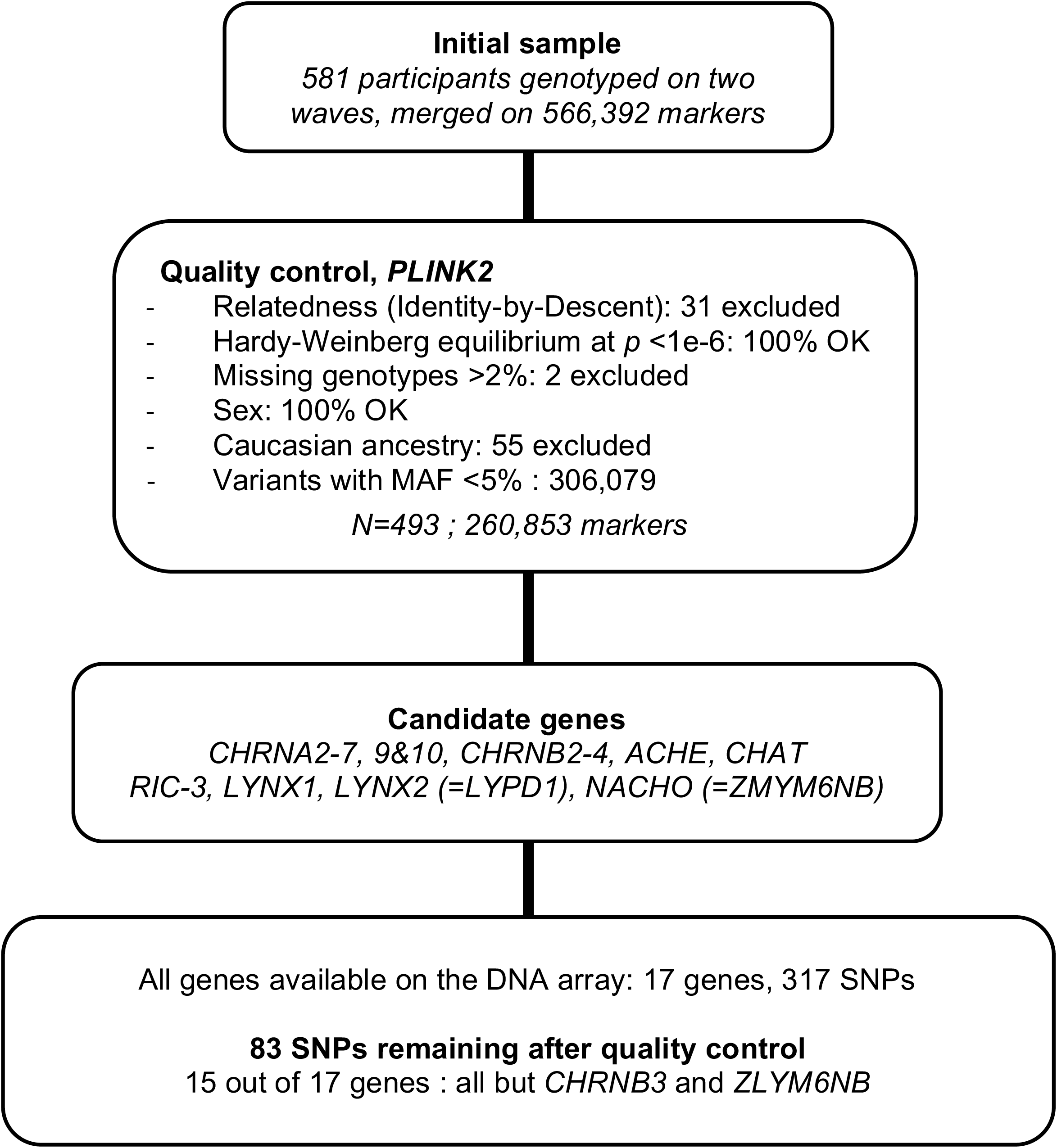
flowchart of the study with participants and variants selection.

Variants showing significant associations with any of the phenotypes of interest were annotated for their biological function and plausible impact according to multiple knowledge-based repositories online, as previously suggested [19]: the *Combined Annotation-Dependent Depletion (CADD)* database [20], brain expression and methylation quantitative trait loci (eQTL through GTEx Analysis Release V8, dbGaP Accession phs000424.v8.p2, https://gtexportal.org/, and mQTL through the mQTLdb database, http://www.mqtldb.org), and their ability to bind DNA enzymes and/or modify DNA conformation (regulomeDB database http://www.regulomedb.org).

### Statistical analyses (Supplementary Table 2)

Single SNP-based tests were first performed with 1/ the number of CPD in current smokers as the dependent variable, by linear regression after squared-root transformation, 2/ current nicotine dependence (HSI score ≥4) as the dependent variable with binary logistic regression (PLINK *--glm* function), 3/ AAO of 1^st^ and 4/ AAO of daily cigarette smoking, by the Chi^2^ log-rank test applied to a Kaplan-Meier survival analysis (R packages *survival* and *survminer*). Hazard ratios (HR) were obtained by cox proportional modelling, after verifying assumption of proportionality for each SNP tested. Gene-based tests for the same phenotypes were then conducted by using either *PLINK ad hoc* function [(*--assoc* or *–logistic*) *perm set-test*] (CPD and HSI data) or the *CoxKM* function (cox model for survival data) from the *KMgene R* package [21] (survival data). Finally, after identification of haplotypes with *HAPLOVIEW* [22] using *PLINK* pedigree files, two locus gene-gene interactions were tested at the SNP level using multidimension reduction methods: *PLINK --epistasis* modifier and general linear model multidimension reduction (GMDR) [12] for CPD and HSI analyses (allele-based tests) and *Cox UM-MDR R* function for survival data [23]. Variants in high LD (r^2^ >0.6) were excluded, so as to consider interactions that would not be due to allele co-segregation, leaving 1431/3403 interactions to be tested.

Variables associated with a given phenotype at Bonferroni-corrected *p* <0.05 were entered as covariates in separate multivariate models to test how they might moderate crude significant genetic associations. In these models, variables with a variance inflation factor (VIF) >2.5 (indicating multicollinearity) were excluded. In case proportionality of HR was not met, time-related variables were split after visual inspection [24] and analysed using the *survSplit* function of R *survival* package. Finally, we computed Bayesian statistics (R packages ‘*BayesFactor*’ and ‘*spBayesSurv*’) in order to (i) ascertain negative findings when corrected *p*-values <0.2 (or when the second most significant association *p*-value was <1) by calculating Bayes Factors (BFs) and (ii) estimate pleiotropy by computing posterior probabilities of associations with all tested phenotypes for each SNP associated with a given phenotype [25]. Strength of Bayesian associations depend on BF values.

Analyses were conducted with PLINK 1.9/2 and R 3.5.3 through R studio 1.1.463 [26] under Mac OS X.12.6 [27]. Statistical significance was set at *p* <0.05 after Bonferroni correction for multiple testing: clinical/sociodemographic variables α =0.05/4 =0.0125 for four smoking phenotypes, SNP-based α =0.05/83 =0.0006, SNP × SNP α =0.05/1431 =0.000024, gene-based α =0.05/number of gene sets valid for testing, and adjusted analyses α =0.05/total number of models that were built across the four phenotypes. R session summary is available in the Supplementary File 1. The present study follows the *STREGA* guidelines for the report of genetic studies [28].

## Results

### Sample description (Supplementary Table 3)

The 493 participants were aged 39 +/- 9 years and 78% were men. There were seven never smokers (missing data =30) and 433 current smokers, who smoked 18 +/- 11 CPD on average (range 1-80) with 39% nicotine dependence according to HSI scores. Smoking began at 14 +/- 3 years, daily smoking at 16 +/-4 years. Shapiro-Wilk tests indicated deviation from normality for CPD and both AAO variables (*p*-values ≤1.7e-15; see Supplementary Figure 2).

### Clinical correlates of smoking phenotypes (Supplementary Table 2)

The number of CPD was significantly associated with lifetime alcohol and sedative use disorders, total number of medications, current treatment for mood disorder. Age at smoking initiation was significantly lower in participants diagnosed with three lifetime SUD or more (polySUD) and with opiate, cannabis and sedative use disorders. AAO of daily smoking was significantly lower in case of polySUD and of cannabis or sedative use disorders; also in case of homelessness.

### Single SNP associations (Table 1, Figures 2&3, Supplementary Figure 3)

**Table 1:**
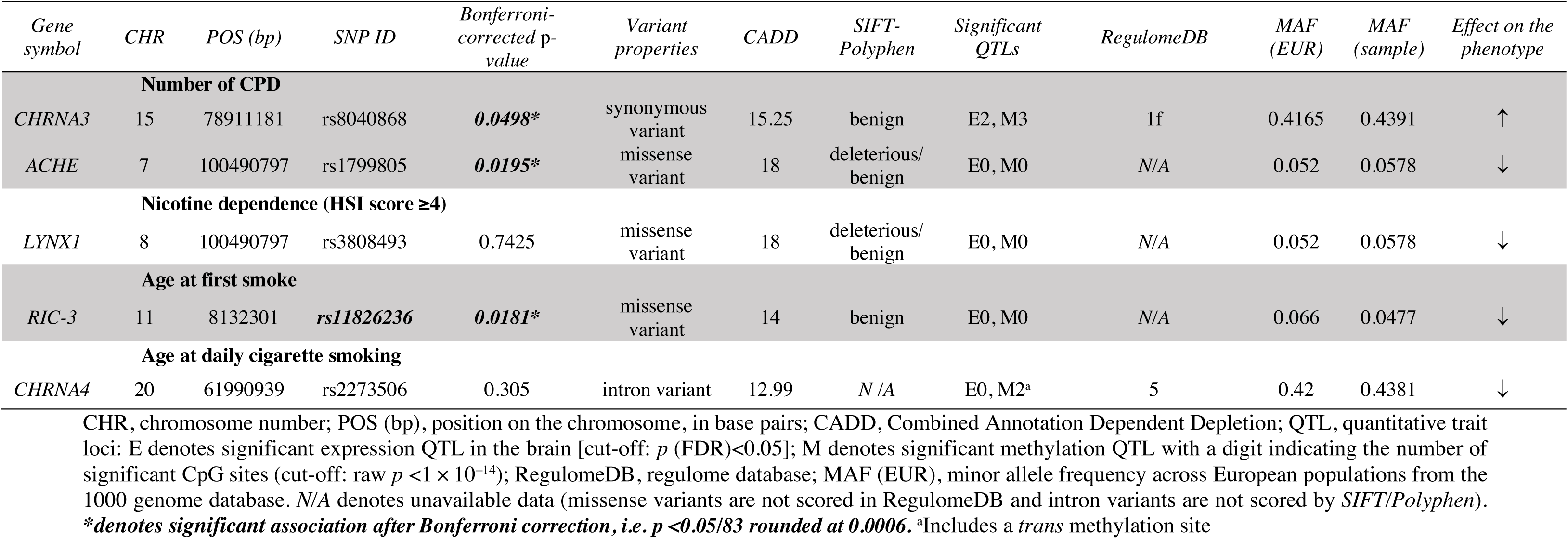
top genetic associations of individual SNP with smoking phenotypes in outpatients with multiple substance use disorders.

After Bonferroni correction, a missense variant in *ACHE*, rs1799805 was associated with a lower number of CPD and a synonymous variant in *CHRNA3*, rs8040868, was significantly associated with a higher number of CPD (Table 1A, Figure 2A). Both associations remained after adjustment for multiple confounders (Figure 2B), with β =−0.76 (SD =0.19, *p* =9.9×10^−5^) for rs1799805 and β =0.34 (SD =0.13, *p* =0.009) for rs8040868; under the recessive model. Lifetime alcohol and sedative use disorders were also independently associated with the number of CPD (β =0.32, SD =0.15, *p* =0.0029 and β =0.31, SD =0.13, *p* =0.0029, respectively) (Figure 2B). Regarding CPD, two SNPs (rs12914385 and rs6495314) had Bonferroni-corrected *p* <0.2. Their BFs were 1.22 and 1.07, respectively, which did not support association with CPD. A missense variant in *RIC3*, rs11826236, was significantly associated with a lower age at smoking initiation (Figure 3A), which remained after multiple adjustments (HR =1.45, *p* =0.028; Figure 3C). Of note, age at smoking initiation had to be split due to non-proportionality of HR (Figure 3B), as described above [24]. Cannabis use disorder was also significantly associated with a lower age at first smoke (HR =1.85, *p* =6.5 × 10^−6^). No association was evidenced regarding AAO of daily cigarette smoking (top SNP =rs2273506 in *CHRNA4*, raw *p* = 0.0037; BF =14, suggesting possible false negative) nor regarding HSI score (top SNP =rs3808493 in *LYNX1*, raw *p* =0.0089; BF =0.78). Summary statistics for all association are in Supplementary Table 4 (Supplementary File 2).

**Figure 2:**
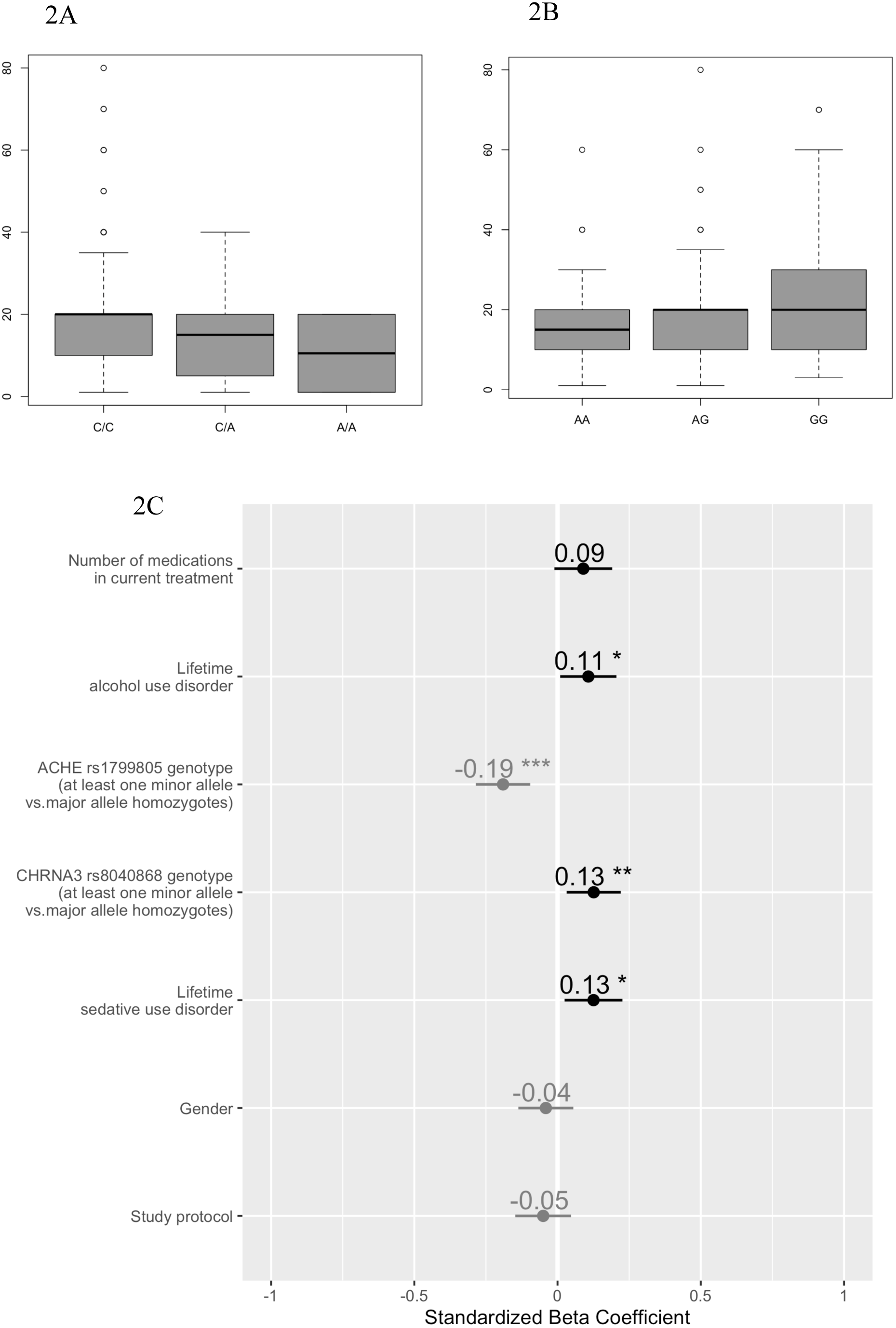
A. Boxplot of the number of cigarettes per day (CPD) according to *ACHE* rs1799805 genotype B. Boxplot of the number of CPD according to *CHRNA3* rs8040868 genotype C. Forest plot of adjusted coefficients (odds ratios) between *ACHE* rs1799805 and *CHRNA3* rs8040868 genotypes and the number of CPD, under the recessive model. * *p* <0.05, \*\**p* <0.01, \*\*\**p* <0.001 (raw values). For each predictor, the line represents the 95% confidence interval.

**Figure 3:**
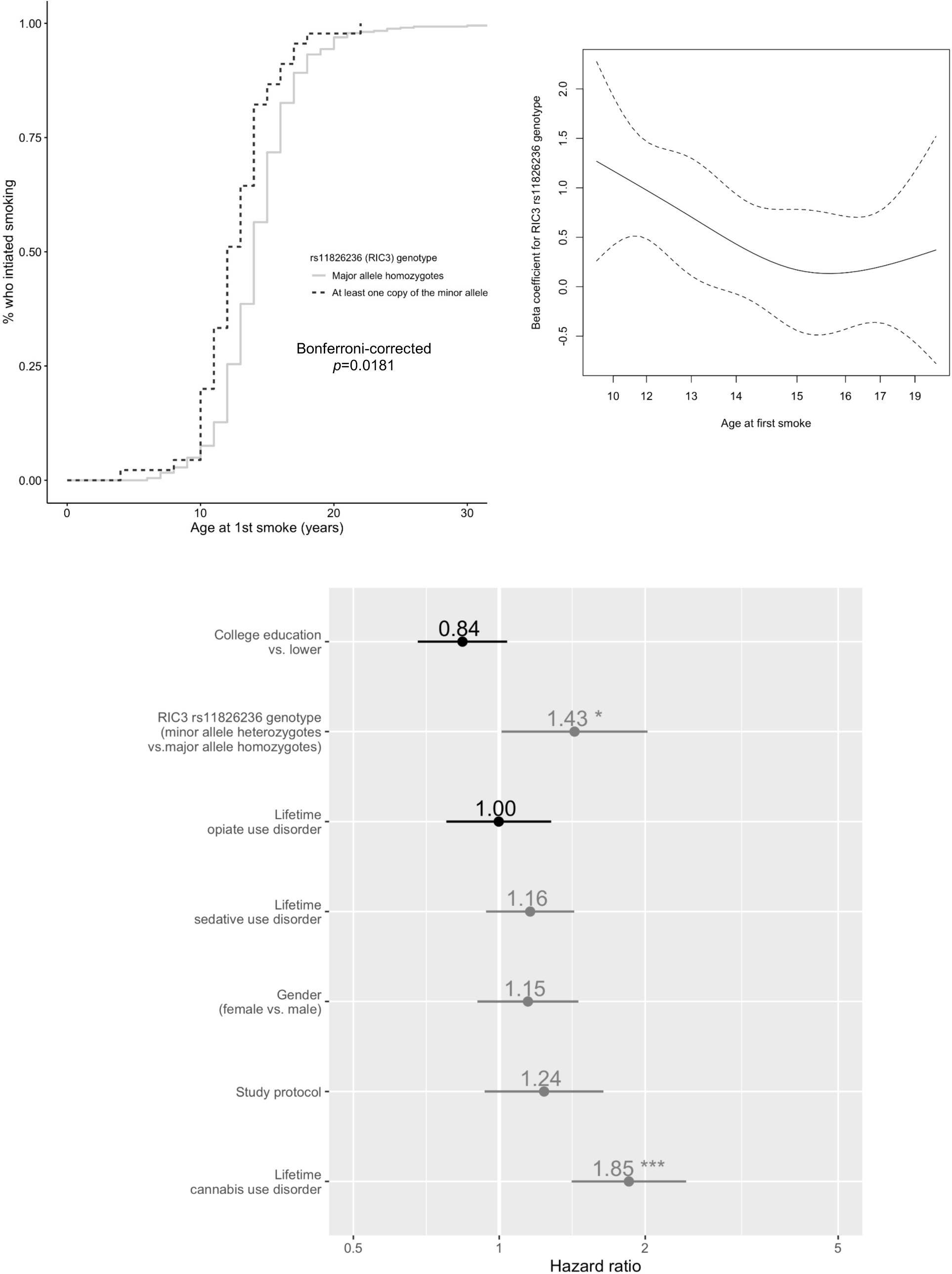
A. Kaplan-Meier curve of the age at 1^st^ smoke as a function of *RIC3* rs11826236 genotype under the recessive model B. Beta coefficient of the Hazard Ratio (HR) of *RIC3* rs11826236 genotype according to age at first smoke, suggesting the non-proportionality of HR D. Forest plot of adjusted HRs for initiating smoking according to *RIC3* rs11826236 genotype (time to smoking initiation split at 13.5 years based on visual inspection of Fig. 3B, due do non-proportionality of HR verified by *ad hoc* statistical testing), under the recessive model. \**p* <0.05, \*\**p* <0.01, \*\*\**p* <0.001 (raw values). For each predictor, the line represents the 95% confidence interval.

### Functional analysis of single SNP associations according to knowledge-based online repositories

GTEx [29], mQTL [30], regulomeDB [31] *and* ENSEMBL (http://grch37.ensembl.org/Homo_sapiens/Info/Index) *(Table 1)* rs1799805 is a missense variant of the acetylcholinesterase gene, with a CADD score of 18, classified as ‘benign’ or ‘deleterious/tolerated’ by the *ENSEMBL* variant effect predictor. It is not associated with *ACHE* expression or methylation. The nAChR α3 subunit gene intron variant rs8040868 is significantly associated with 19 eQTLs, including a lower expression of *CHRNA5* in the cortex (normalized effect size, NES =−0.65) and increased expression of *CHRNA5* antisense RNA RP11-650L12.2 in the basal ganglia (NES =−0.51). It is also associated with increased methylation on three *cis* sites, suggesting additional negative effect on *CHRNA3* expression. Moreover, this locus has a high affinity for both transcription factors and DNA-enzymes (regulomeDB score =1f).

rs11826236 is a missense variant in the ‘resistance to inhibitors of cholinesterase 3’ gene, a chaperone protein for nAChRs. This SNP has a CADD score of 14, classified as ‘benign’ by the *ENSEMBL* variant effect predictor. It is significantly associated with 6 eQTLS, mostly related to decreased *RIC3* expression in non-neural peripheral tissue.

### Gene-based tests

Gene-based tests confirmed the association between the number of CPD and *ACHE* (Bonferroni-*corrected p* <0.05 after 46614 permutations), which resisted multiple adjustment (*p* =0.0002. See Figure 2C for adjustment variables) and was driven by rs1799805 only. There was no gene-based association with HSI score (lowest Bonferroni-corrected *p* =0.2467 for *LYNX1*). *RIC3* was associated with lower age at smoking initiation (Bonferroni-corrected *p* = 0.0166), which was lost after multiple adjustment (Bonferroni-corrected *p* =1). *CHRNA4* was associated with a lower AAO of daily tobacco smoking (Bonferroni-corrected *p* =0.0475), which remained after multiple adjustment.

### Multi-SNP analyses: haplotypes and gene × gene interactions (Table 2)

The variants that were tested in the present study belonged to 17 haplotype blocks (Supplementary Figure 5), none of which encompassed the significant SNPs displayed in Table 1. There was no SNP × SNP interaction regarding CPD, HSI score nor age at smoking initiation with either PLINK or MDR methods. Conversely, two SNP × SNP interactions were evidenced regarding AAO of daily tobacco smoking. They involved *CHAT* rs10776585 × *CHRNA4* rs6090392 (HR =1.68, corrected *p* =0.0388) and *RIC3* rs4758042 × *CHRNB4* rs11072793 (HR =1.99, corrected *p* =0.0167), both remaining after multiple adjustments (HR =1.69, corrected *p* =0.0001 and HR =1.85, corrected *p* =0.0004, respectively).

**Table 2:**
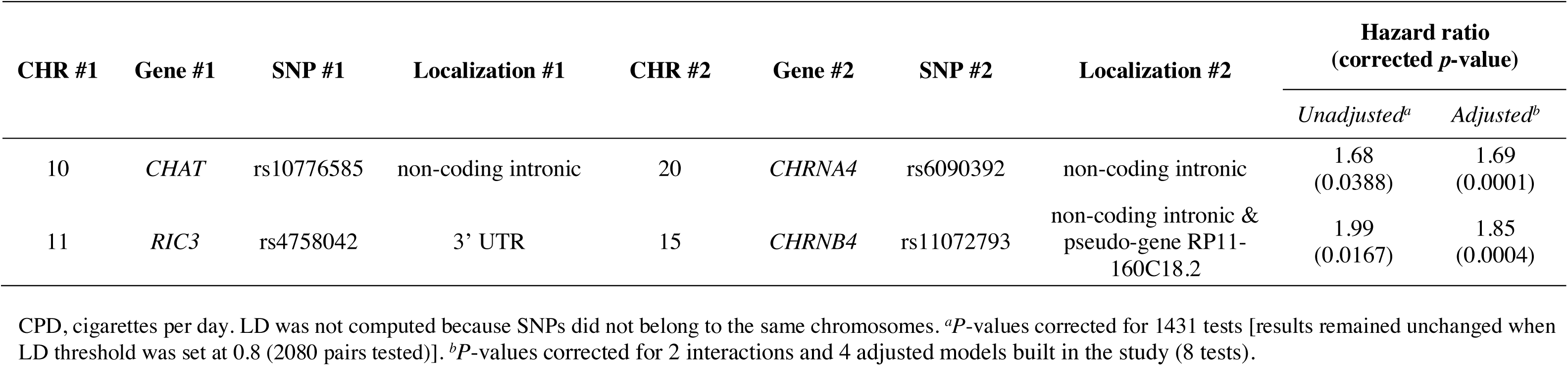
SNP × SNP interactions in the AAO of daily tobacco smoking according to the *coxumMDR* dimension reduction function. *P*-values were obtained after 1000 permutations and Bonferroni correction for 3403 tests.

### Multiple phenotypes analysis: pleiotropy (Supplementary Figure 4)

BFs ranged from 0 (no association) for *CHRNA4* rs2273506 to 21(very strong association) for *CHRNA3* rs8040868. For three of the seven top SNP with BF suggesting at least some true association (see *methods* section and summary statistics in Supplementary File 2), posterior probabilities indicated shared associations across smoking phenotypes, especially between CPD and AAO of daily smoking, CPD and HIS, and CPD and both AAOs.

## Discussion

Patients with multiple SUDs attending specialized care are characterized by high levels of daily tobacco smoking (90%), nicotine dependence (39% of current smokers) (sample data) and addictive comorbidity. To our knowledge, this is one of the first published studies to address these issues altogether in a pathway-wide genetic study. The four main genetic findings were: *CHRNA3* rs8040868 associated with a higher number of CPD, *ACHE* rs1799805 associated with a lower number of CPD, *RIC3* rs11826236 associated with a lower age at smoking initiation, two SNP × SNP interactions involved in AAO of daily smoking, and pleiotropy for rs8030868 and rs1799805 regarding all smoking phenotypes.

The present study replicated the previously identified associations between *CHRNA3* and smoking behaviours, including the number of CPD, which typically involved rs1051730 [14], a plausible tag marker for functional haplotypes in the *CHRNA5-A3-B4* cluster. This makes rs8040868 a relatively new SNP of interest in the cluster. This SNP is in strong LD (r^2^ >0.8) with 25 SNPs on chromosome 15, including *CHRNA5* rs16969968, which has been repeatedly associated with heavy tobacco smoking [6]. According to *GTEx portal*, rs8040868 is associated with decreased *CHRNA5* expression in the brain. It is also in strong LD with rs146009840, which is associated with decreased expression of a *CHRNA5* antisense *RNA (*RP11-650L12.2) in the caudate nucleus. The co-inheritance of these three SNPs could thus 1/ reduce *CHRNA5* expression, which has been associated with a lower risk of nicotine dependence [32] and 2/ have both enhancing and lowering effects on the receptor function through the decrease of RP11-650L12.2 expression (which limits *CHRNA5* protein translation) and rs16969968, respectively. Of note, the apparent discrepancy between results regarding HSI *vs.* CPD may be consistent with the present findings. In fact, nicotine dependence assessed by the HSI yielded no genetic association in the present sample, suggesting that the genetic influence of nAChRs may rather exert on the number of CPD than on other symptoms of loss of control over nicotine intake. In this population, both phenomena may thus rely on more distinct risk factors than in the general population, suggesting that tobacco smoking may further be used to cope with other drugs *craving*/adverse effects [33] or attention deficits [34], which may overall be mediated by the relationship of these symptoms with perceived stress [35]. The discrepancy between CPD and HSI has been previously reported [36], and HSI itself remains a proxy for the full DSM nicotine dependence syndrome. Our pleiotropy analysis indirectly supported this finding. Thus, *CHRNA3* rs8040868 showed particularly strong evidence for shared associations across smoking phenotypes (BF =21, see [25]), mainly between CPD and AAO of daily tobacco smoking, and *ACHE* rs1799805 showed opposite directions in its posteriors between CPD and HIS (Supplementary Figure 4).

To our knowledge, this is the first published evidence of association between *RIC3* and smoking behaviour. The *RIC3* gene as a whole and the missense mutation rs11826236 have been previously associated with cognitive decline [37–39], and this SNP is in strong LD (r^2^ >0.8) with 795 SNPs on chromosome 8; none of which, however, elicits major structural or regulatory impact. *RIC3* is crucial to the trafficking of homomeric nAChRs (α7)5 and of serotonin 3A receptors (5-HT3A) [40]. The gene activity overall enhances target receptor expression at the cell surface [40]. This may be related to age at 1^st^ smoking by modulating cognitive dimensions involved in addictive processes such as procedural learning through (α7)_5_ nAChRs [41,42] and impulsivity through 5-HT3Rs [43]. The phenotype ‘age at 1^st^ smoke’ is of utmost relevance since up to 69% adolescents who initiate smoking will be daily smokers at some point [44]. Adolescent brains may be particularly susceptible to the psychopathological effects of tobacco smoke [45].

Gene-based analyses were in line with SNP-based findings and, interestingly, also yielded an association between *CHRNA4* and earlier AAO of daily smoking. This is in line with previous findings of associations between *CHRNA4* SNPs and AAO of nicotine dependence [46].

Two SNP × SNP interactions were associated with a lower AAO of daily smoking after multiple adjustments. From a molecular perspective, none reflected nAChR subunit arrangement. The first involved two non-coding introns from *CHAT* and *CHRNA4* with low affinity for DNA transcription factor/enzymes (both regulomeDB scores =4). *CHAT* rs10776585 is associated with increased *AGAP6* gene expression in the cerebellum (*AGAP6* encodes a protein that functions as a putative GTPase-activating protein in the cell nucleus). The second interaction between *RIC3* and *CHRNB4* involved 3’-UTR and pseudo-gene introns with low affinity for DNA transcription factor/enzymes (regulomeDB score =6 for rs4758042 and unavailable for rs11072793). *RIC3* rs4758042 is associated with reduced cerebellar expression of *TUB*, which encodes the signal transduction and DNA-binding protein *Tubby protein homolog*, and of *RIC3* (normalized effect sizes −0.29 to −0.34). *CHRNB4* rs11072793 also maps to the downstream pseudogene *RP11-160C18.2* and has been associated with earlier regular smoking initiation within a 5-order SNP interaction pattern in male subjects of Korean ancestry [47]. Interestingly, this interaction involved two other SNPs from *RP11-160C18.2*, strengthening its potential role in smoking initiation phenotypes. Moreover, rs11072793 is associated with increased expression of *ADAMTS7* in the brain cortex (normalized effect size =0.32), which encodes a metalloprotease that was previously associated with body mass index in smokers [48]. From a broader neurobiological perspective, the interaction between *CHRNA4* and *CHAT* may reflect a more global link between the role of the synthesis of acetylcholine and high-affinity nAChRs such as α4β2-containing, and help clarifying the role of *CHRNA4* suggested by other findings (see our own gene-based tests and [46]) while the interaction between *RIC3* and *CHRNB4* is likely related to the role of *Ric3* in the proper folding and assembly of β4 subunits (https://www.uniprot.org/uniprot/P30926#interaction) and to the multiple regulatory roles of *RP11-160C18.2*. Overall, the pattern of gene × gene interactions we evidenced in the AAO of daily smoking is supported by the biological function of the SNPs that are involved – considered both individually and under their interaction. It also highlighted gene expression regulation in the cerebellum, the role of which is increasingly recognized in addiction, to nicotine [49,50]. No significant interaction was found for the other smoking phenotypes, suggesting that more systems/circuits are progressively recruited in the transition from smoking initiation toward more complex – truly representative of the state of addiction – phenotypes [6]. Of note, both AAO phenotypes involved *RIC3* variants, which may reflect its association with an endophenotype associated with both the motivation to initiate and maintain tobacco smoking.

Limitations must be acknowledged. Some possibly relevant variants were absent from the DNA array, notably rs880395 and rs1948, which modulate *CHRNA5* and *CHRNB4* expression, respectively [7]. There was up to 11% missing values regarding HSI scores, but this lack was randomly distributed in the core phenotypic variables (data not shown) and was thus unlikely to have biased the results. We did not record the proportions of proposed/eligible/included participants. Among computed Bayes Factors (BFs)s, only one was strongly suggestive of a false negative result between *CHRNA4* rs2273506 and AAO of daily tobacco smoking (corrected *p* =0.3, BF =14). Finally, we only studied pairwise gene × gene interactions. With that regard, we believed the modest sample size and the relatively large number of markers that were investigated would have prevented the study robustness and interpretability in case higher-order interactions would have been tested.

The present study has several strengths. It relied on standard and validated procedures for genetic QC, functional assessment of identified variants by using multiple databases, and phenotyping. Phenotypic characterization was extensive, allowing for studying complementary smoking phenotypes, other associated phenotypes and possible confounders in multivariate models, including for survival data. The smoking phenotypes tested here capture a large proportion of the complex phenomenology of tobacco smoking/nicotine addiction while maintaining the transfer potential of our results into preclinical models. A significant number of high-quality genetic markers of relevant effectors in the nAChR system, including metabolism/chaperone/trafficking proteins that were recently characterized, were tested in single SNP, gene-based and gene × gene analyses, relevant to the pathophysiology of complex disorders. Minor allele frequencies in the sample were close to those obtained from 1000G samples, further supporting the absence of population stratification. Finally, there were no biases due to missing data (data not shown).

The sample reflected populations with multiple and rather severe SUD and might not generalize to people with less severe disorders, e.g. with a single SUD, drawn from general population samples. Importantly, in France, specialized care settings for SUDs are organized to remain as accessible as primary care centers to treatment-seeking individuals, even if they are hospital-based, and they must warrant both anonymity and free access (https://solidarites-sante.gouv.fr/IMG/pdf/08_79t0.pdf).

To conclude, the present study provides multiple-level, yet preliminary evidence that, in people at high-risk for severe smoking outcomes, gene variants from the nicotinic pathway may further moderate the risk for specific tobacco smoking phenotypes. Associations showed both a polygenic and a pleiotropic nature that included SNP × SNP interactions, in line with previous research in less specific populations [14]. Taken altogether, our findings also highlight the importance of metabolic and chaperone/trafficking proteins in the pathogenicity of nAChRs. Such associations warrant further biochemical research to better understand their functional implications, especially as regards the less investigated genes in tobacco smoking such as *CHAT, ACHE, RIC3* and *CHRNA4*.

## Supporting information

SNP list from the nicotinic pathway

## Acknowledgements

The authors would like to thank for his precious help in implementing the *coxUM-MDR* function in our study and the investigators that were involved in building the present cohort. Gaël Dupuy (Assistance Publique – Hôpitaux de Paris, CSAPA Murger, Hôpital Fernand Widal, Paris, F75010), Didier Touzeau (CSAPA Clinique liberté, Bagneux F-92220), Cyrille Orizet (Assistance Publique – Hôpitaux de Paris, CSAPA Montecristo, Hôpital Européen Georges Pompidou, Paris F-75015), Philippe Coeuru (CSAPA Espoir Goutte d’Or – Aurore, Paris F-75018), Pierre Polomeni (Assistance Publique – Hôpitaux de Paris, Service d’addictologie, Hôpital René Muret-Bigottini, Sevran F-93270), Xavier Laqueille (Centre Hospitalier Sainte-Anne, CSAPA Moreau de Tours, Paris F-75014), Elisabeth Avril (CSAPA, Gaia association, Paris F-75011), Anne-Marie Simonpoli (Assistance Publique – Hôpitaux de Paris, ELSA, Hôpital Louis Mourier, Colombes, F-92700, Pr. Olivier Cottencin (Service d’Addictologie, Hôpital Fontan 2 – CHRU Lille, Lille F-59037), Belforte (Assistance Publique – Hôpitaux de Paris, CSAPA Montecristo, Hôpital Européen Georges Pompidou, Paris F-75015), Aurélia Gay (CHU Saint-Etienne, Pôle Psychiatrie Adultes et Infanto-Juvénile, Saint-Etienne F-42055), Philippe Lack (CSAPA Hôpital de la Croix-Rousse, Lyon F-69317), Philippe Coeuru (CSAPA Espoir Goutte d’Or – Aurore, Paris, F-75018). Philippe Batel (Assistance Publique – Hôpitaux de Paris, Unité de Traitement Ambulatoire des Maladies Addictives, Hôpital Beaujon, Clichy F-92110), Philippe Batel (Clinique Montevideo, Boulogne F-92100), Jean-Baptiste Trabut (Hôpital Emile Roux, Limeil-Brevannes, F-94450).

This study makes use of data generated by the Wellcome Trust Case-Control Consortium. A full list of the investigators who contributed to the generation of the data is available from www.wtccc.org.uk. Funding or the project was provided by the Wellcome Trust under award 076113, 085475 and 090355” and cite the relevant primary WTCCC publication (details of which can be found on the WTCCC website).

## Funding

- *Investissements d’Avenir* program managed by the ‘Agence Nationale de la Recherche’ (ANR) under reference ANR-11-IDEX-0004-02 and Labex BIO-PSY;
- DRCI (OST07013) and French Ministry of Health (PHRC National 2010 AOM10165) for patients recruitment;
- ANR (ANR-13-SAMA-0005-01), ERA-net Neuron *Synapse and Mental Disorders* 2013 (COCACE, R14026KK) and 2017 (ADIKHUMICE, ANR-17-NEU3-0002-05) for clinical and genetic analysis of the combined sample
-This study was designed and conducted during a ‘Poste d’accueil’ research fellowship obtained by Dr Romain ICICK from Assistance Publique – Hôpitaux de Paris and Labex BIO-PSY.

---

**Supplementary Table 1 (supplementary file 2): original (before quality control, see Figure 1) list of 314 markers from the nicotinic pathway, ordered by chromosome number and position.**

**Supplementary Table 2:**
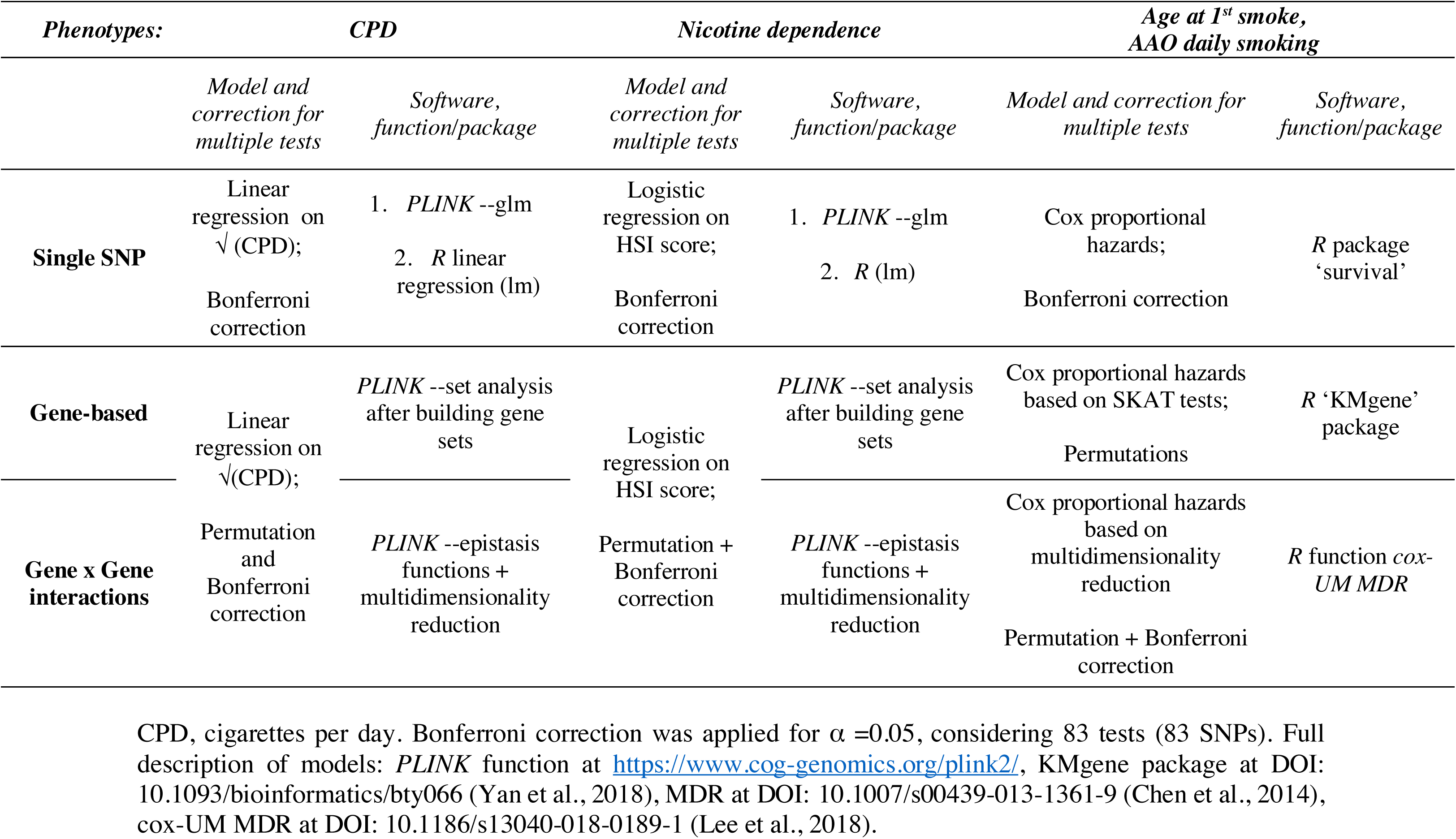
software, software packages and related statistical models and correction methods for multiple testing used in the study

**Supplementary Table 3:**
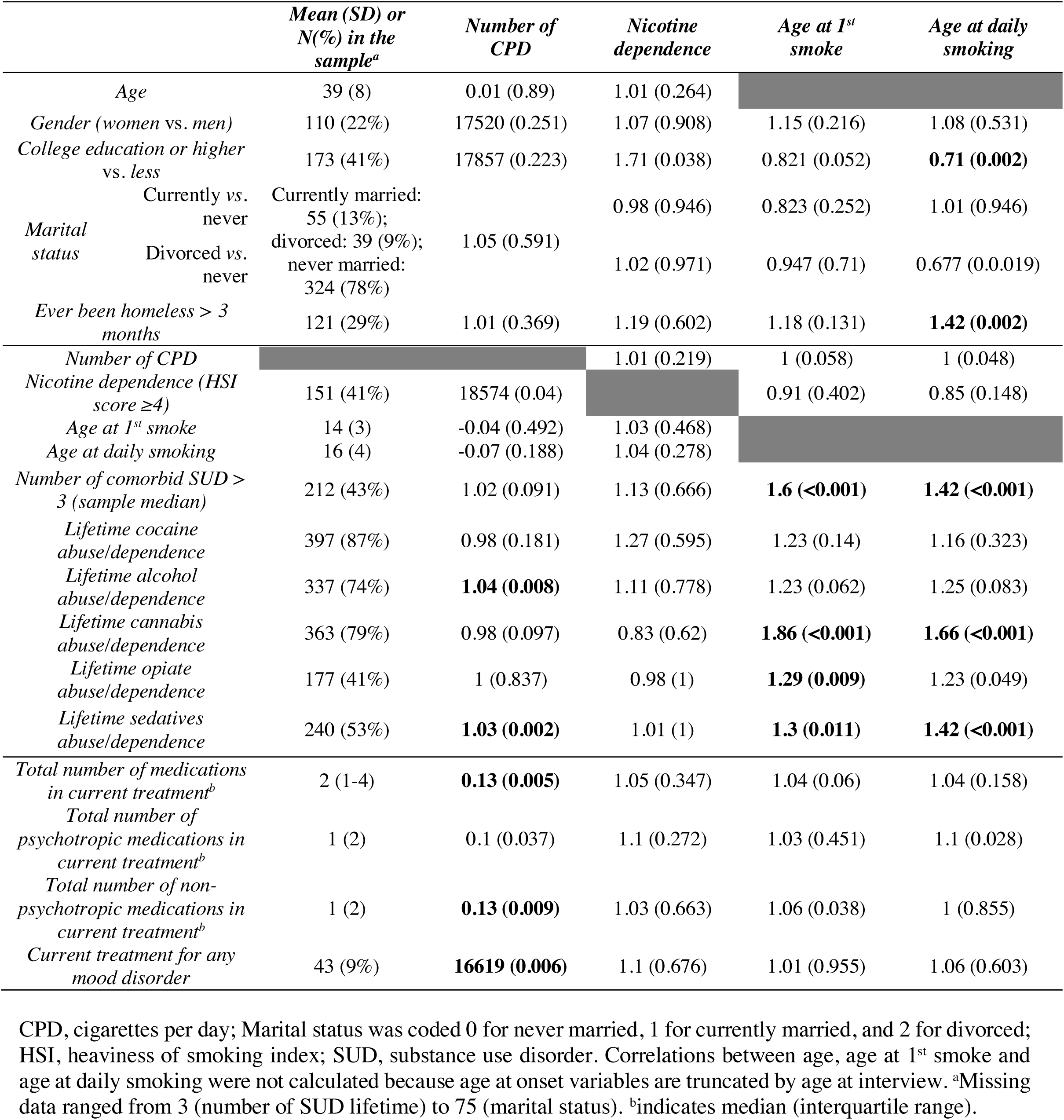
associations between the number of CPD, nicotine dependence, age at first smoke and age at daily smoking with clinical and sociodemographic variables in the sample of outpatients seeking treatment for SUDs (N =493). Chi-squared for categorical × categorical, Wilcoxon for binary × continuous, Kruskal-Wallis for non-binary categorical, Spearman for continuous × continuous, and cox proportional hazards for survival data were used. Unadjusted univariate odds ratios, correlation coefficients or hazard ratios are displayed, rounded to the 2^nd^ decimal, with their *p*-values, rounded to the 3^rd^ decimal. Values are in bold for *p*-values <0.05 after Bonferroni correction for four tests.

**Supplementary Table 4: (supplementary file 2): summary statistics for associations between 83 SNPs of the nicotinic pathway and the squared roots of the number of cigarettes smoked per day (CPD), nicotine dependence (HSI), age at first smoke (TBAGE1ST) and AAO daily tobacco smoking (TBDAILY).**

**Supplementary Figure 1:**
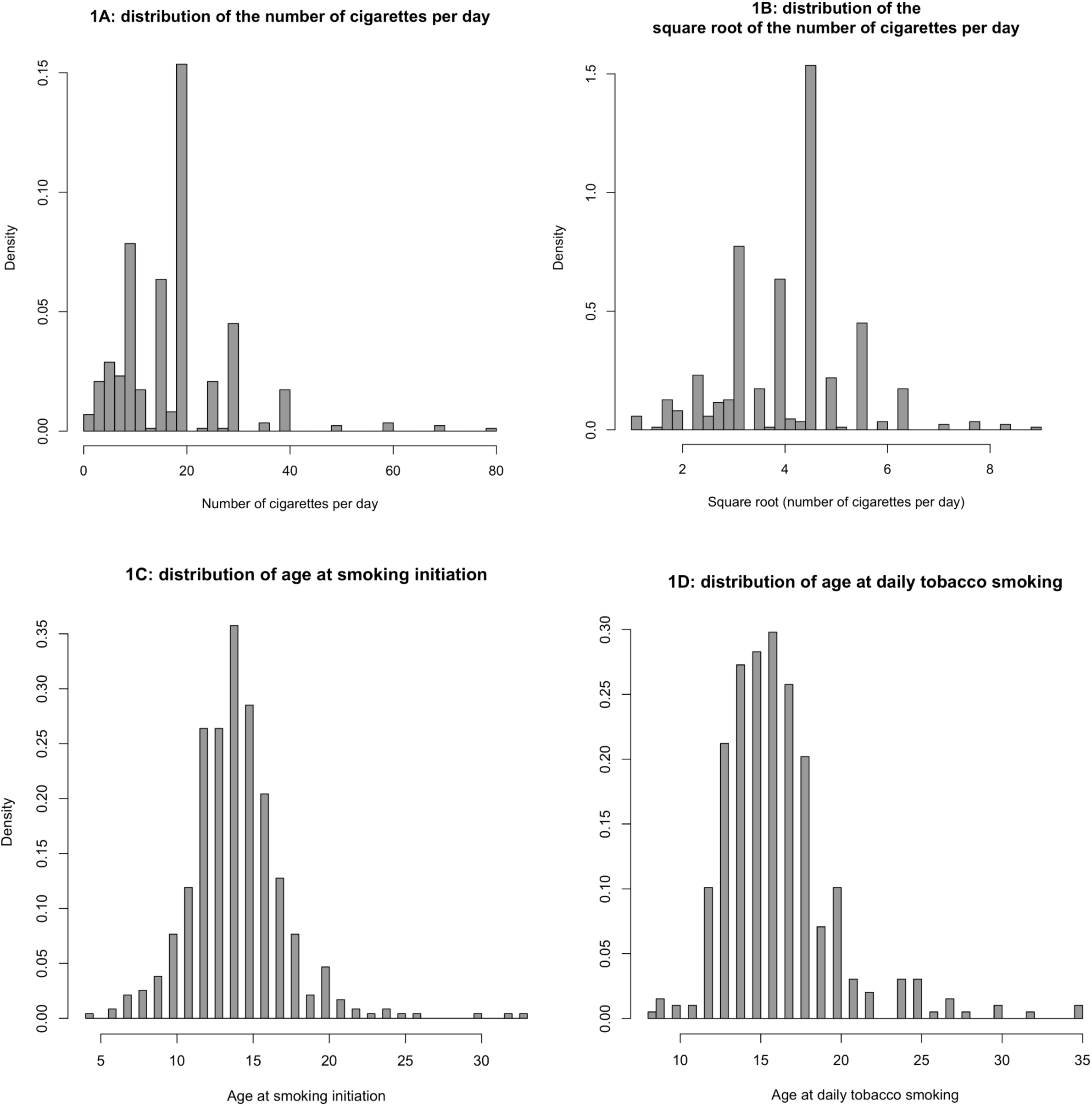
distributions of the (A) number of cigarettes per day (CPD, N =433), (B) CPD after square root transformation, (C) age at smoking initiation (N =470) and (D) age at daily smoking (N =396) in the sample.

**Supplementary Figure 2:**
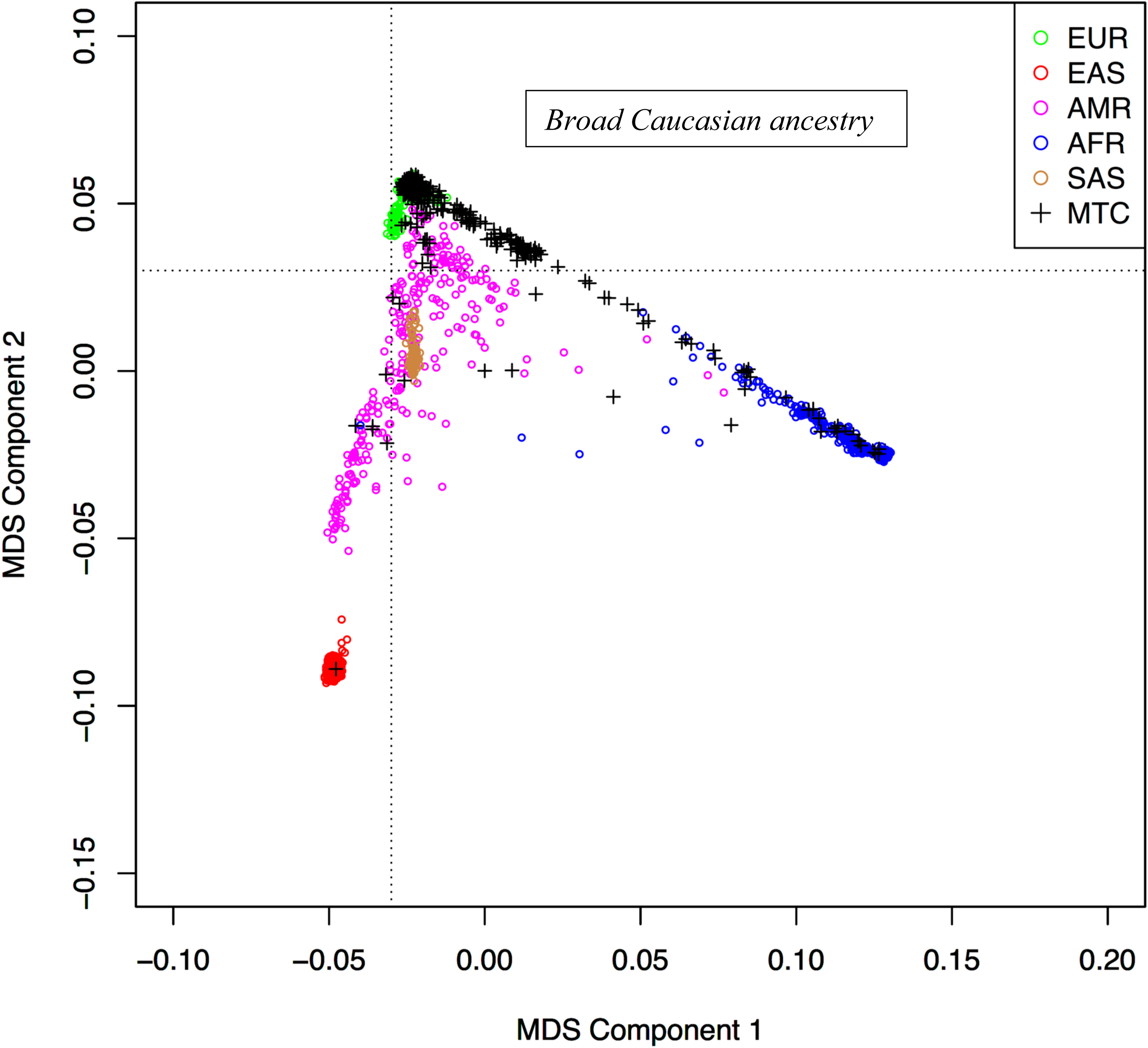
principal component analysis of the study sample and 1000G, phase 3 superpopulations. Dotted lines indicate cut-offs for considering Caucasian ancestry. MDS, multidimensional scale; EUR, Europe; EAS, Eastern Asia; AMR, America; AFR, Africa; SAS, South America; MTC, study sample.

**Supplementary figure 3:**
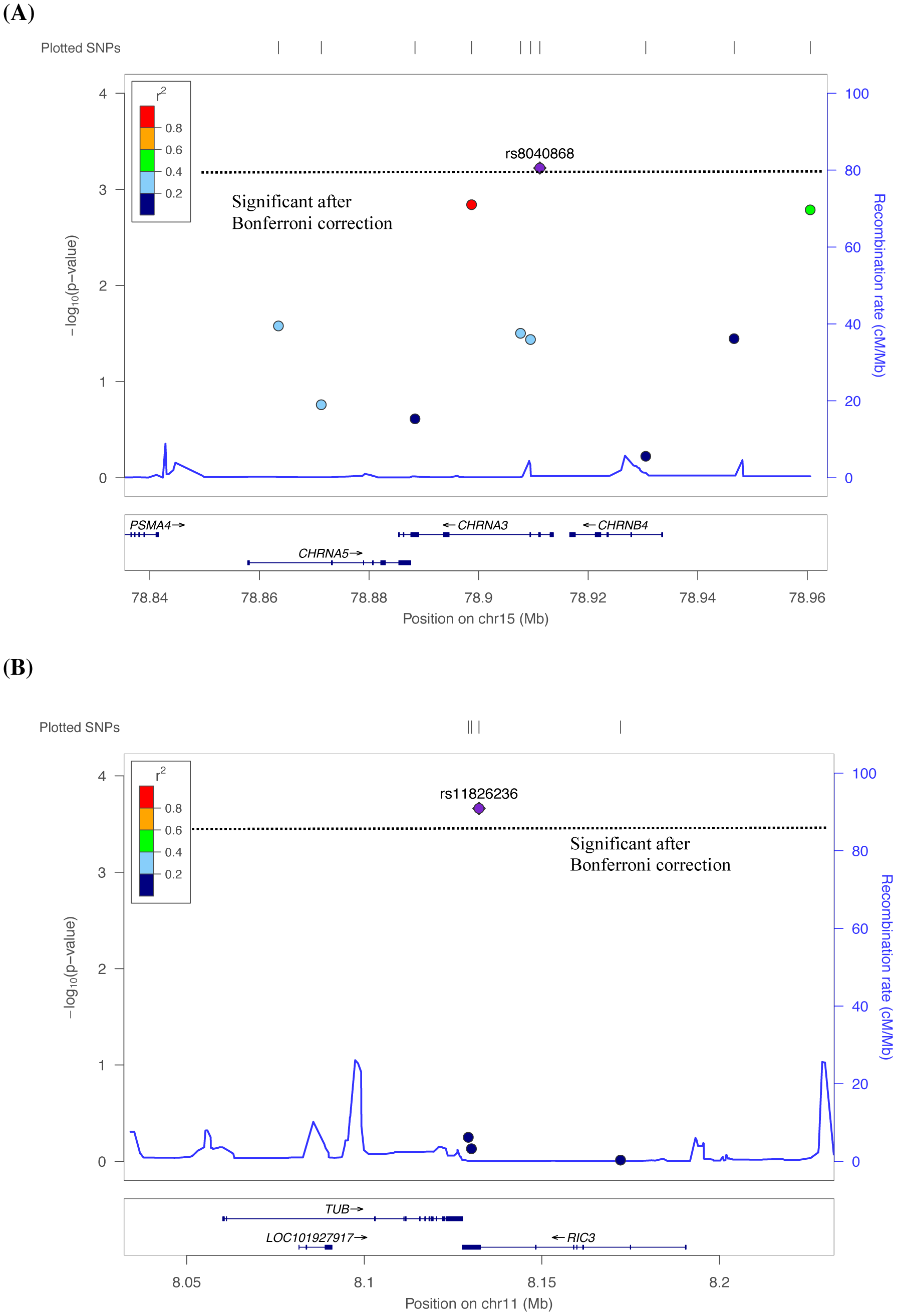
regional plot (obtained at http://locuszoom.org/genform.php?type=yourdata) of *p*-values in (A) the *CHRNA3-CHRNA5-CHRNB4* cluster tested with the squared root of the number of cigarettes per day (linear regression) and (B) *RIC3* tested with age at 1^st^ smoke (cox proportional hazards). The dotted line indicate significance cut-off after Bonferroni correction (*p* =0.05/83 rounded at 0.0006). (A)

**Supplementary Figure 4:**
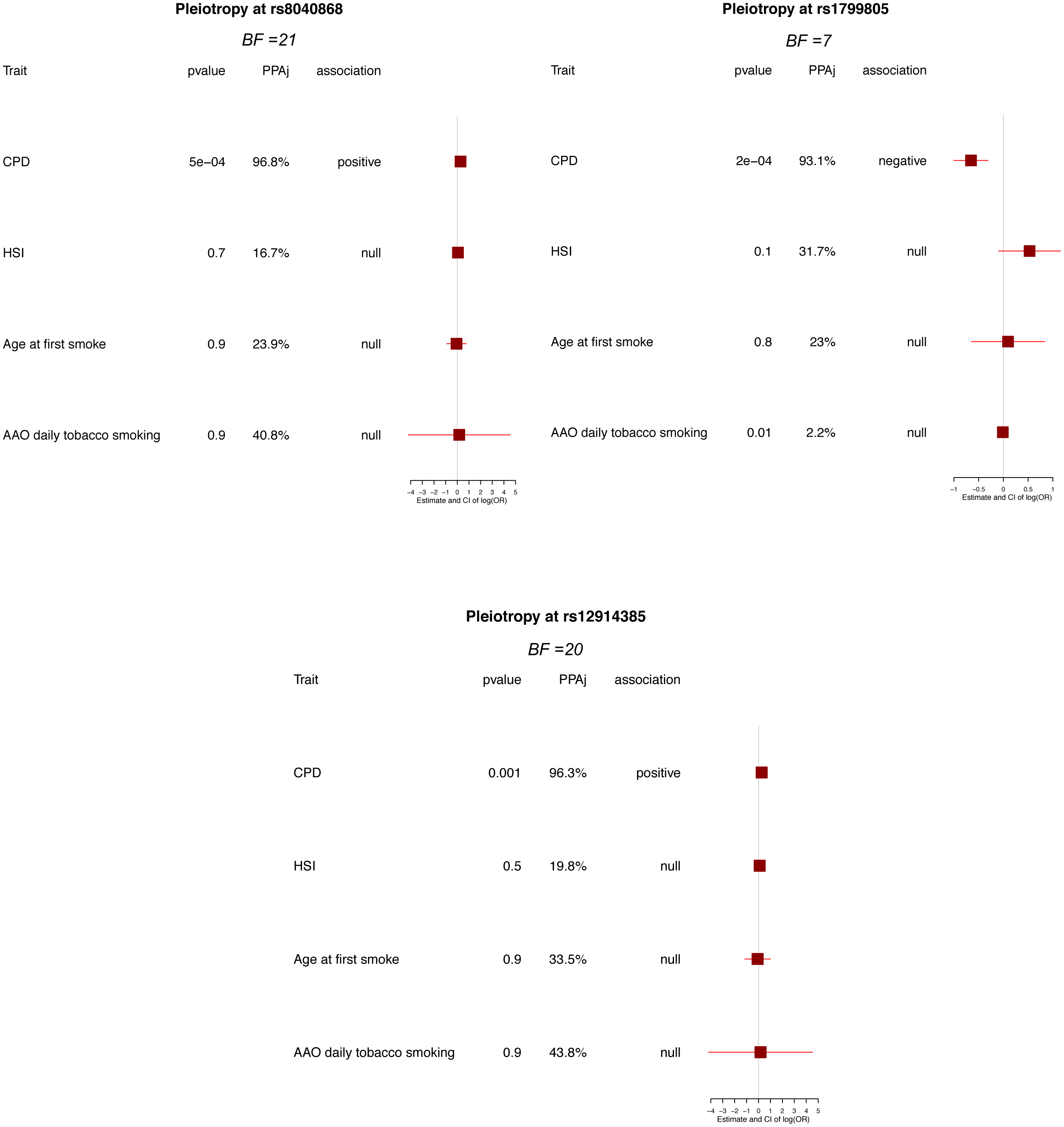
evidence of pleiotropy regarding three of the seven top associations evidenced in the study (BFs >3, suggestive of association, were considered for displaying here). Forest plots were obtained with the *CPBayes* package for *R* after 10,000 iterations and indicate the posterior probability of association for each trait and global Bayes Factor (BF), along with the strength and significance of the initial association. CPD, cigarettes per day; HSI, heaviness of smoking index; AAO, age at onset.

**Supplementary Figure 5:**
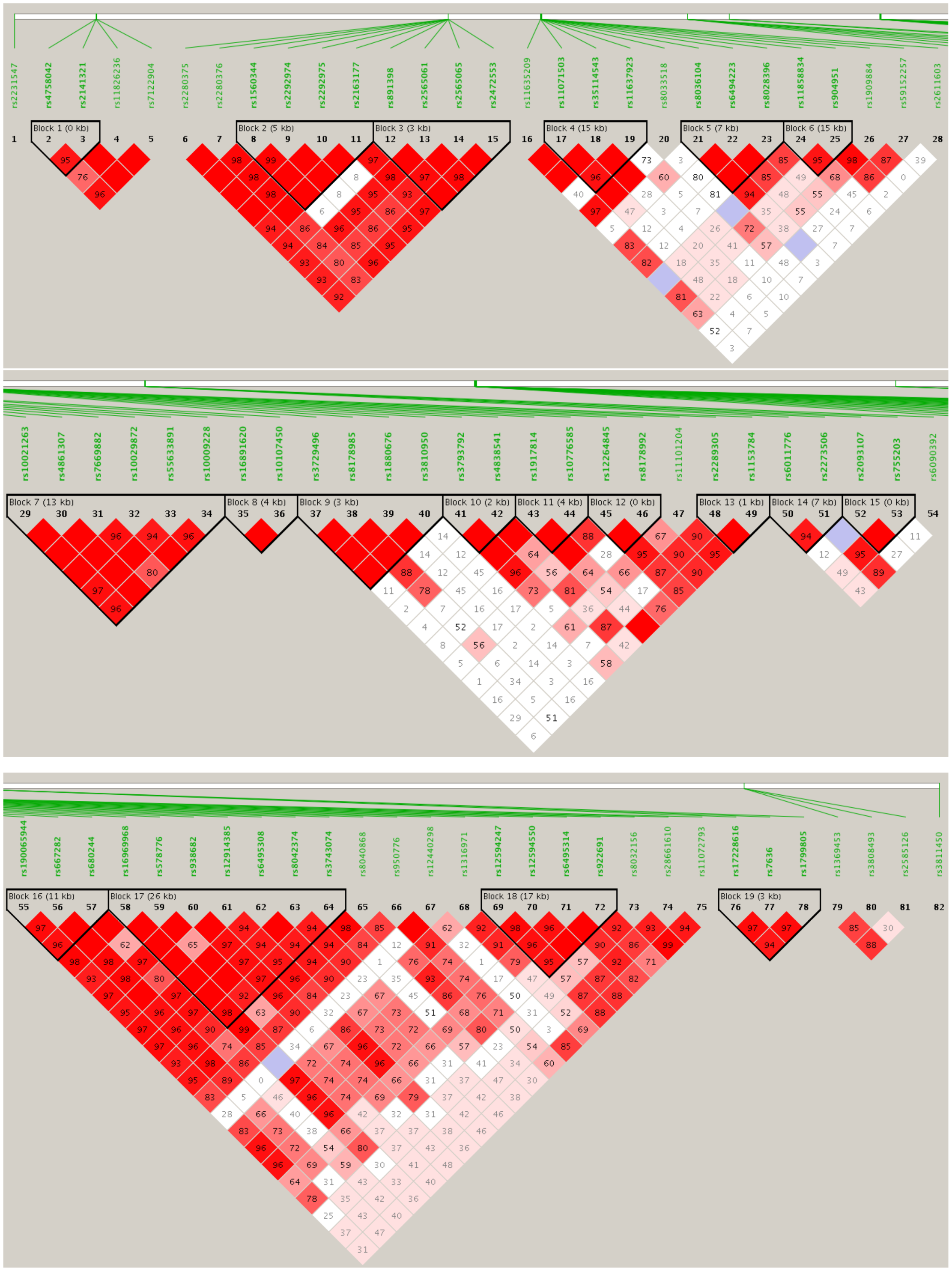
LD plot of the 17 haplotype blocks identified by *HAPLOVIEW* (figure was truncated in three contiguous parts following blocks order along chromosomes)

