## Supplementary methods file for "Gene-Gene interactions and pleiotropy in the brain nicotinic pathway associated with the heaviness and precocity of tobacco smoking among outpatients with multiple substance use disorders"

1. *Sample recruitment*

Treatment-seeking outpatients attending tertiary care programs in the Paris area, mostly around the 'Gare du Nord' train station, were recruited through two multicentric research protocols (Mouly et al., 2015a; Vorspan et al., 2015a) aimed at deciphering the phenotypic and genotypic architecture of severe SUDs (Icick et al., 2017). Both protocols were approved by the local ethics committees (CPP Ile-de-France VI for study one and CPP Ile de France IV for study two; NCT00894452 and NCT01569347, respectively). Participants were French-speaking, 18+ years old individuals, seeking treatment in any of the participating centers. Further inclusion criteria were:

- Study one (S1, seven sites) = receiving stable methadone treatment for three months or more for treating opiate dependence [see (Mouly et al., 2015b) for details on study protocol];
- Study two (S2, six sites) = any past-year cocaine use [see (Vorspan et al., 2015b) for details on study protocol].

For the present study, participants were excluded if they were undergoing compulsory treatment or were unable to consent for any other reason (non-French speaking, major cognitive impairment). The level of comorbidity was not considered for exclusion.

2. *Clinical assessments*

The data were obtained during a single interview conducted by trained psychologists or M.D.s.

- Sociodemographic conditions were collected using a standard procedure, and included the number of years in education starting from elementary school, marital status, and history of homelessness, which was retained if the participant reported having spent at least three months living on the streets;
- Lifetime patterns of use were characterized for each substance through the E section of the Structured Clinical Interview for DSM (SCID) for DSM-IV (First et al., 1996), comprising diagnoses of SUD and age at onset (AAO) of substance use and of SUD.

- Participants' ongoing medication was collected through free text during the interview. The total number of medication was computed, and the total number of psychotropic medication was computed separately. For this, we used the international Anatomical Therapeutic Chemical (ATC) classification, codes N06A (antidepressants) and N06BA (centrally acting sympathomimetic psychostimulants). ATC codes N05CA (barbiturates), N05CD08 (midazolam) and N05CM05 (scopolamine) were excluded because these molecules are not used for psychiatric indications. ATC code R06AD01 (the widely used hypnotic alimemazine) was searched for manually because it is classified under the respiratory system group. Mood-stabilizers, as authorized in France as of March, 19th 2019 (HAS, 2017), were extracted manually because they belonged to different ATC groups.

- Investigators used standard survey tips to determine these events with the best possible accuracy (Kendig et al., 2014).

3. *References*

First, M., Spitzer, R., Gibbon, M., Williams, J., 1996. Structured Clinical Interview for DSM Disorders [WWW Document]. URL http://www.scid4.org/faq/scidfaq.html (accessed 2.29.16).

HAS, 2017. Haute Autorité de Santé - ALD n° 23 - Troubles bipolaires [WWW Document]. URL https://www.has-sante.fr/portail/jcms/c_849818/fr/ald-n-23-troubles-bipolaires (accessed 3.19.19).

Icick, R., Karsinti, E., Lépine, J.-P., Bloch, V., Brousse, G., Bellivier, F., Vorspan, F., 2017. Serious suicide attempts in outpatients with multiple substance use disorders. Drug Alcohol Depend 181, 63–70. https://doi.org/10.1016/j.drugalcdep.2017.08.037

Kendig, H., Byles, J.E., O'Loughlin, K., Nazroo, J.Y., Mishra, G., Noone, J., Loh, V., Forder, P.M., 2014. Adapting data collection methods in the Australian Life Histories and Health Survey: a retrospective life course study. BMJ Open 4, e004476. https://doi.org/10.1136/bmjopen-2013-004476

Mouly, S., Bloch, V., Peoc'h, K., Houze, P., Labat, L., Ksouda, K., Simoneau, G., Declèves, X., Bergmann, J.F., Scherrmann, J.-M., Laplanche, J.-L., Lepine, J.-P., Vorspan, F., 2015a. Methadone dose in heroin-dependent patients: role of clinical factors, comedications, genetic polymorphisms and enzyme activity. Br J Clin Pharmacol 79, 967–977. https://doi.org/10.1111/bcp.12576

Mouly, S., Bloch, V., Peoc'h, K., Houze, P., Labat, L., Ksouda, K., Simoneau, G., Declèves, X., Bergmann, J.F., Scherrmann, J.-M., Laplanche, J.-L., Lepine, J.-P., Vorspan, F., 2015b. Methadone dose in heroin-dependent patients: role of clinical factors, comedications, genetic polymorphisms and enzyme activity. Br J Clin Pharmacol 79, 967–977. https://doi.org/10.1111/bcp.12576

Vorspan, F., Fortias, M., Zerdazi, E., Karsinti, E., Bloch, V., Lépine, J.-P., Bellivier, F., Brousse, G., van den Brink, W., Derks, E.M., 2015a. Self-reported cue-induced physical symptoms of craving as an indicator of cocaine dependence. Am J Addict 24, 740–743. https://doi.org/10.1111/ajad.12303
Vorspan, F., Fortias, M., Zerdazi, E.-H., Karsinti, E., Bloch, V., Lépine, J.-P., Bellivier, F., Brousse, G., van den Brink, W., Derks, E.M., 2015b. Self-reported cue-induced physical symptoms of craving as an indicator of cocaine dependence. Am J Addict 24, 740–743. https://doi.org/10.1111/ajad.12303

4. R session summary obtained with *sessionInfo*

R version 3.5.3 (2019-03-11)
Platform: x86_64-apple-darwin15.6.0 (64-bit)
Running under: macOS Sierra 10.12.6

Matrix products: default
BLAS:
/System/Library/Frameworks/Accelerate.framework/Versions/A/Frameworks/vecLib.framework/Versions/A/libBLAS.dylib
LAPACK: /Library/Frameworks/R.framework/Versions/3.5/Resources/lib/libRlapack.dylib

locale:
[1] en_US.UTF-8/en_US.UTF-8/en_US.UTF-8/C/en_US.UTF-8/en_US.UTF-8

attached base packages:
[1] parallel  stats4    stats     graphics  grDevices utils     datasets  methods   base

other attached packages:
 [1] reshape2_1.4.3       doBy_4.6-2           plyr_1.8.4           pipeR_0.6.1.3
 [5] GenomicRanges_1.34.0 GenomeInfoDb_1.18.2  IRanges_2.16.0       S4Vectors_0.20.1
 [9] BiocGenerics_0.28.0  qqman_0.1.4

loaded via a namespace (and not attached):
 [1] Rcpp_1.0.2             compiler_3.5.3         pillar_1.4.2           XVector_0.22.0
 [5] bitops_1.0-6           tools_3.5.3            zlibbioc_1.28.0        tibble_2.1.3
 [9] lattice_0.20-38        pkgconfig_2.0.2        rlang_0.4.0            Matrix_1.2-17
[13] rstudioapi_0.10        writexl_1.1            yaml_2.2.0             xfun_0.9
[17] GenomeInfoDbData_1.2.0 stringr_1.4.0          dplyr_0.8.3            knitr_1.24
[21] grid_3.5.3             tidyselect_0.2.5       glue_1.3.1             calibrate_1.7.2
[25] R6_2.4.0               purrr_0.3.2            magrittr_1.5           MASS_7.3-51.4
[29] assertthat_0.2.1       stringi_1.4.3          RCurl_1.95-4.12        crayon_1.3.4
